# Wheat pathogen *Zymoseptoria tritici N*-myristoyltransferase inhibitors: on-target antifungal activity and an unusual metabolic defense mechanism

**DOI:** 10.1101/2020.02.09.940536

**Authors:** Roman O. Fedoryshchak, Cory A. Ocasio, Ben Strutton, Jo Mattocks, Andrew J. Corran, Edward W. Tate

## Abstract

*Zymoseptoria tritici* is the causative agent of *Septoria tritici* blotch (STB), which costs billions of dollars annually to major wheat-producing countries in terms of both fungicide use and crop loss. Agricultural pathogenic fungi have acquired resistance to most commercially available fungicide classes, and the rate of discovery and development of new fungicides has stalled, demanding new approaches and insights. Here we investigate a potential mechanism of targeting an important wheat pathogen *Z. tritici* via inhibition of *N*-myristoyltransferase (NMT). We characterize *Z. tritici* NMT biochemically for the first time, profile the *in vivo Z. tritici* myristoylated proteome and identify and validate the first *Z. tritici* NMT inhibitors. Proteomic investigation of the downstream effects of NMT inhibition identified an unusual and novel mechanism of defense against chemical toxicity in *Z. tritici* through the application of comparative bioinformatics to deconvolute function from the previously largely unannotated *Z. tritici* proteome. Research into novel fungicidal modes-of-action is essential to satisfy an urgent unmet need for novel fungicide targets, and we anticipate that this study will serve as a useful proteomics and bioinformatics resource for researchers studying *Z. tritici*.

## INTRODUCTION

*Zymoseptoria tritici* (formerly *Mycosphaerella graminicola*) is a fungal pathogen of wheat causing *Septoria tritici* blotch (STB) in areas of wheat cultivation^1,2^. The blotch usually goes unnoticed in the first 10 days of infection, but later manifests itself as leaf necrosis and leads to rapid plant death in the following 10-20 days^3^, during which time fungal spores maturate and spread through leaf-to-leaf contacts and rain splash dispersion. STB is the fungal disease causing the greatest reduction in wheat yields in Europe^4^ and has a growing footprint in North America. Control of STB relies heavily on scheduled regimens of different fungicides sprayed over the field throughout the year^5^, with STB alone accounting for up to 70% of fungicide usage in the European Union. *Z. tritici* incurs losses of billions of dollars annually for the agriculture industry, in terms of disease control cost as well as reducing potential wheat yields, and current trends indicate a worsening situation due to the emergence and spread of strains of *Z. tritici* resistant to commonly used fungicides^1^.

Fungicides play an important role in maintaining crop yields in wheat cultivation. However, populations of *Z. tritici* have acquired field resistance against two of the four mainstream antifungal classes: firstly, methyl benzimidazole carbamates (MBC)^6^ and later the quinone outside inhibitors (QoI)^7,8^. Of the remaining classes, demethylase inhibitors (DMI) often display decreased performance in the field due to reduced sensitivity of *Z. tritici*^9^, whilst the latest succinate dehydrogenase inhibitors (SDHI) have been effective to date. However, studies under laboratory conditions have shown that *Z. tritici* can acquire resistance as a result of single-point mutations in succinate dehydrogenase^10,11^.

Although it may be possible to genetically engineer crop protection against STB^2^, the most relied upon strategy remains development of new antifungals. *N*-myristoyltransferase (NMT) is a well-characterized, druggable, and essential enzyme which was widely pursued as an antifungal target for human disease in the 1990s and early 2000s. NMT catalyzes co-translational attachment of a myristoyl moiety to the N-terminal amine of proteins carrying an N-terminal glycine followed by a certain consensus sequence. NMT inhibition is effective in targeting a range of pathogens including parasites from *Plasmodium, Leishmania* and *Trypanosoma*^12^ and fungi from *Candida*^13^ and *Cryptococcus*^14^ genera. A few inhibitor scaffolds were reported to target NMT in clinically relevant fungi *Candida albicans, Aspergillus fumigatus* and *Cryptococcus neoformans*^15^, but were so selective in their fungal species spectrum^16^ that they failed to fulfil the broad spectrum antifungal properties required at the time of their discovery. More recently, a series of *Trypanosoma brucei* NMT inhibitors were shown to have limited activity against *A. fumigatus* NMT^17^. Recent validation of NMT as an anti-parasitic^18–20^ and antiviral^21^ target has been accompanied by accumulation of substantial new data encompassing NMT functional networks, inhibitor scaffolds and crystal structures describing structure-activity relationships. These advancements have revitalized interest in NMT as a target and prompted us to consider it in an as yet unexplored agricultural context.

In this paper, we report the first characterization of *Z. tritici* NMT *in vitro* and the identification and characterization of the *Z. tritici N*-myristoylated proteome *in vivo*. We also report the discovery of a new class of fungal NMT inhibitors which are the most potent discovered to date, with low nanomolar potency against the enzyme and analyze their mode of action *in vivo* through whole-proteome response to treatment with inhibitor. Finally, we combine these data to discuss the defense mechanisms employed by *Z. tritici* to counter NMT inhibition, and the opportunities and challenges presented by NMT as a target in STB.

## RESULTS AND DISCUSSION

### Characterization of ZtNMT and identification of highly potent ZtNMT inhibitors

We cloned and expressed *Z. tritici* NMT (ZtNMT) as a construct lacking the first 99 amino acids (ZtNMTΔ99); this construct excludes the unstructured ribosome binding region of NMT which is known in many other organisms to be both dispensable for enzymatic activity and prone to cause problems with expression and purification^22–24^. First, we established a CPM (7-diethylamino-3-(4’-maleimidylphenyl)-4-methylcoumarin) based assay for ZtNMTΔ99 measuring release of free coenzyme A thiol when substrate (Myr-CoA) is processed by the enzyme (Figures 1A and S1A, S1B)^25^. As the peptide substrate we used an octapeptide derived from the *N-*terminus of F9×1I2 (*Z. tritici* predicted homolog of ADP-ribosylation factor 1, ARF1), which we hypothesized would be natively myristoylated *in vivo*.

**Figure 1.**
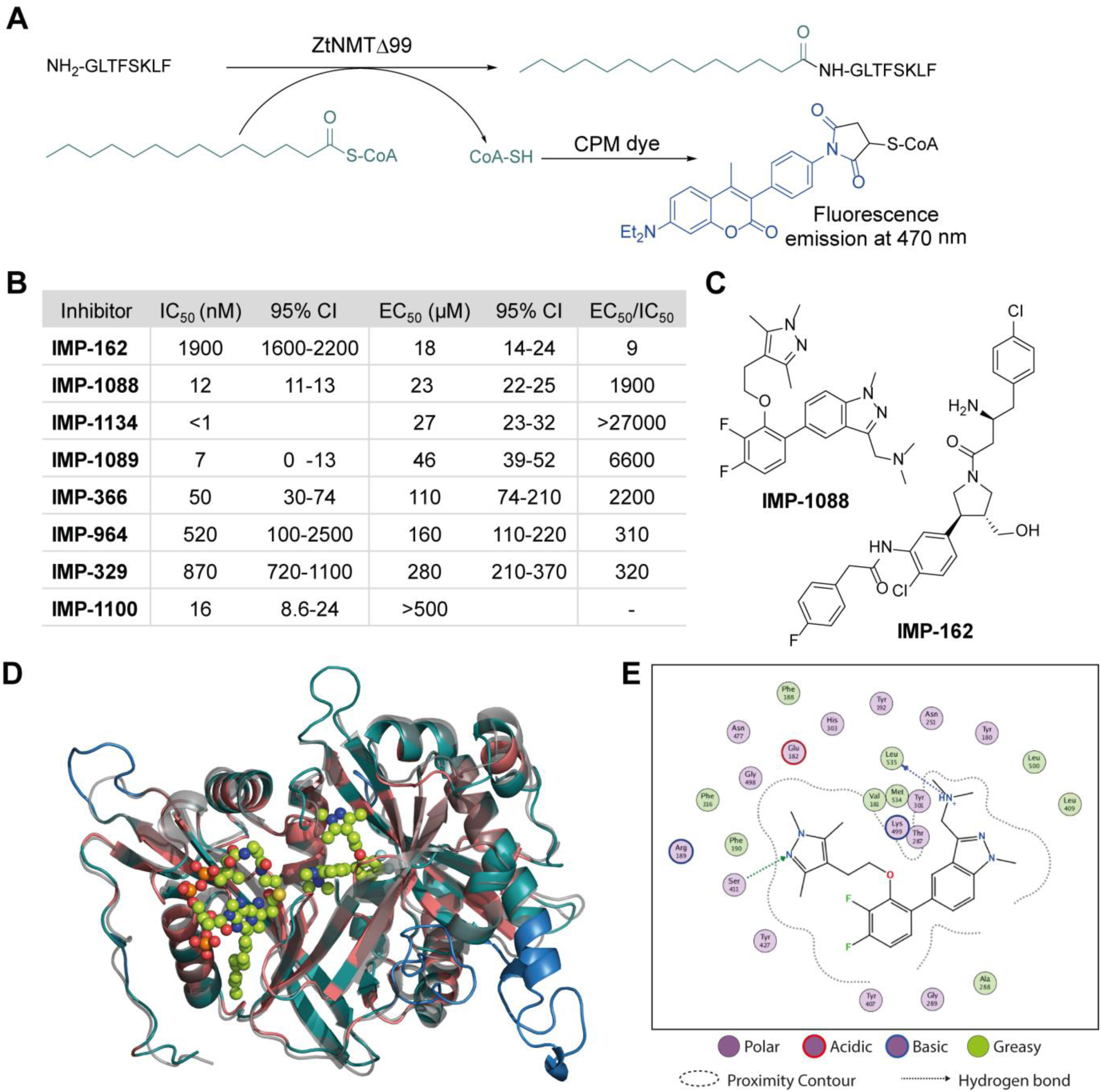
ZtNMT enzymatic activity and structure. **A**) In an *in vitro* assay, ZtNMT transfers myristate from myristoyl-CoA to the N-terminal glycine of a substrate peptide. The CPM reagent enables fluorogenic readout of the reaction rate based on the generation of coenzyme A thiol. **B**) Potency of selected inhibitors in enzymatic (CPM) and *Z. tritici* cell viability assays (**IMP-366** is also known as **DDD85646**^32^). **C**) Inhibitors **IMP-162** and **IMP-1088** were found to be among the most potent in the viability assays. **IMP-162** is an outlier with an apparently better translation from enzyme to *in vivo* inhibition. **D**) ZtNMT homology model generated using Swiss Model^33^ based on a reported *A. fumigatus* NMT X-Ray structure 4UWI^32^ was aligned to HsNMT1 (PDB: 5MU6, in transparent grey). HsNMT1 structure contains small molecules **IMP-1088** (center) and myristoyl-CoA (left) shown in stick-and-ball representation. Structural alignment of the two enzymes indicates the highly conserved fold with a backbone RMSD of 2.259 Å, and identifies identical amino acids (red), non-conserved residues (green), and a predicted helix extensions in ZtNMT (blue). **E**) A map of ligand interactions between docked **IMP-1088** and its induced fit generated ZtNMT receptor protein residues.

As a starting point to identify potential ZtNMT inhibitors, we used a set of available inhibitors of different structural series originally developed against NMTs from various unrelated species. Most inhibitors in our set displayed at least some activity, with several highly potent examples identified in the low nanomolar IC_50_ range (Figures 1B and 1C; inhibitor structures and full assay data are provided in Supplemental Table S1). We next sought to understand the enzyme-inhibitor interactions which influence activity. Our attempts to crystallize ZtNMT resulted in slow-growing and poorly diffracting microcrystals (not shown). Therefore, we constructed a homology model of ZtNMT (Figure 1D) and explored inhibitor binding by alignment to known experimental crystal structures. Whilst ZtNMT shares the highest similarity to other fungal NMTs (e.g. overall 40% identity with *C. albicans* NMT and 47% with *A. fumigatus*, Figure S1C), it also has >30% sequence identity with human NMT1 (HsNMT1; unlike unicellular eukaryotes, higher eukaryotes possess at least two NMT paralogues, NMT1 and NMT2). To understand inhibitor interactions with ZtNMT, we aligned a ZtNMT homology model based on the closely related *Aspergillus fumigatus* NMT to a structure of HsNMT1 bound to inhibitor **IMP-1088** (Figure 1C)^21^.

The peptide binding pocket, which is where most NMT inhibitors are known to bind, clearly shows conservation of aromatic and hydrophobic residues binding **IMP-1088** (Figure 1D), including ZtNMT Ser411 (Ser405 in HsNMT1) which forms a conserved hydrogen bond to the pyrazole in **IMP-1088. IMP-1088** also appears to have the same interactions with ZtNMT *C-*terminal carboxyl of Leu535 residue (Gln496 in HsNMT1). The C*-*terminal acid of Leu535 is critical for enzymatic activity since it plays a role in coordination of the protein substrate N-terminus and forms a strong ionic interaction with the amine of the inhibitor. Whilst **IMP-1088** is an ultrapotent inhibitor of HsNMT1 with *K*_d_ estimated <200 pM, it is >60-fold less potent against ZtNMT (12 nM IC_50_).

We next undertook docking studies to establish structure activity relationship of **IMP-1088** interactions with ZtNMT. To generate a homology model suitable for docking, we energy minimized **IMP-1088** overlaid from the HsNMT1 structure in our ZtNMT model. The atomic coordinates of **IMP-1088** were used to set the binding site, enabling induced fit docking of a subset of **IMP-1088** derivatives using MOE (see Supporting Methods). The docked inhibitors were scored based on S-score value, root mean square deviation (RMSD) between poses and similarity to the **IMP-1088** binding mode in the ZtNMT homology model (Table S1). The induced fit docking simulation yielded binding free energies ranging from −7.93 to −9.46 kcal/mol, covering about 2 orders of magnitude in binding affinity, and overall had very consistent binding poses with RMSD values ranging from 0.820 – 1.61 Å (Table S1). To assess quality of docking, we compared the **IMP-1088** docked pose with our **IMP-1088** energy minimized ZtNMT homology model. Docked **IMP-1088** maintained key interactions between Ser411 and the Leu535 backbone and both models eliminated steric clash between the fluorinated central ring and Tyr301 (Figures 1E and S2B). The SAR generated using these docking parameters revealed two key scaffolds, triazolopyridine and imidazopyridine, with maximal binding energy when the pyrazole ring of the R_1_ group was substituted with a tertiary amide and the R_2_ group substituted on the core scaffold was a tertiary or secondary amine (Figure S2A). To substantiate this docking study, we also overlaid the top four hits, containing these structural features, with docked **IMP-1088** in the ZtNMT homology model, revealing similar binding modes (Figure S2B). We also validated the predictive value of this study as the second ranked hit, **IMP-1134**, displayed the greatest biochemical potency against ZtNMT (IC_50_ < 0.001 μM, Table S1).

Anti-proliferative activity (EC_50_) for several most potent compounds was measured over 40 hours using a MTS-based assay, a colorimetric assay that tracks the metabolism of a tetrazolium-based MTS dye in living cells^26^, revealing a strong drop-off in activity *in vivo* (Figure 1B). **IMP-1088** and **IMP-162** (Figure 1C) displayed EC_50_ around 20 μM in this assay, and whilst a few other indazole structures had IC_50_ below 100 μM, the remainder were inactive. A similar drop-off in activity from enzyme to in-cell inhibition (of 1000-fold or greater) has been seen previously for NMT inhibitors in the context of *Leishmania*^27^ and the fungus *Aspergillus fumigatus*^17^, but its origin has not been explored in detail. As evidenced by our data, the IC_50_ to EC_50_ drop-off in activity is not limited to any structural series of inhibitors but appears to be wide-ranging, and may be due to interference in uptake, active efflux, or metabolic degradation. **IMP-162** is a marked outlier in our measurements since it shows little activity against ZtNMT *in vivo* (see below, Figure 2C), suggesting that the mode of action of this compound is unrelated to NMT.

**Figure 2.**
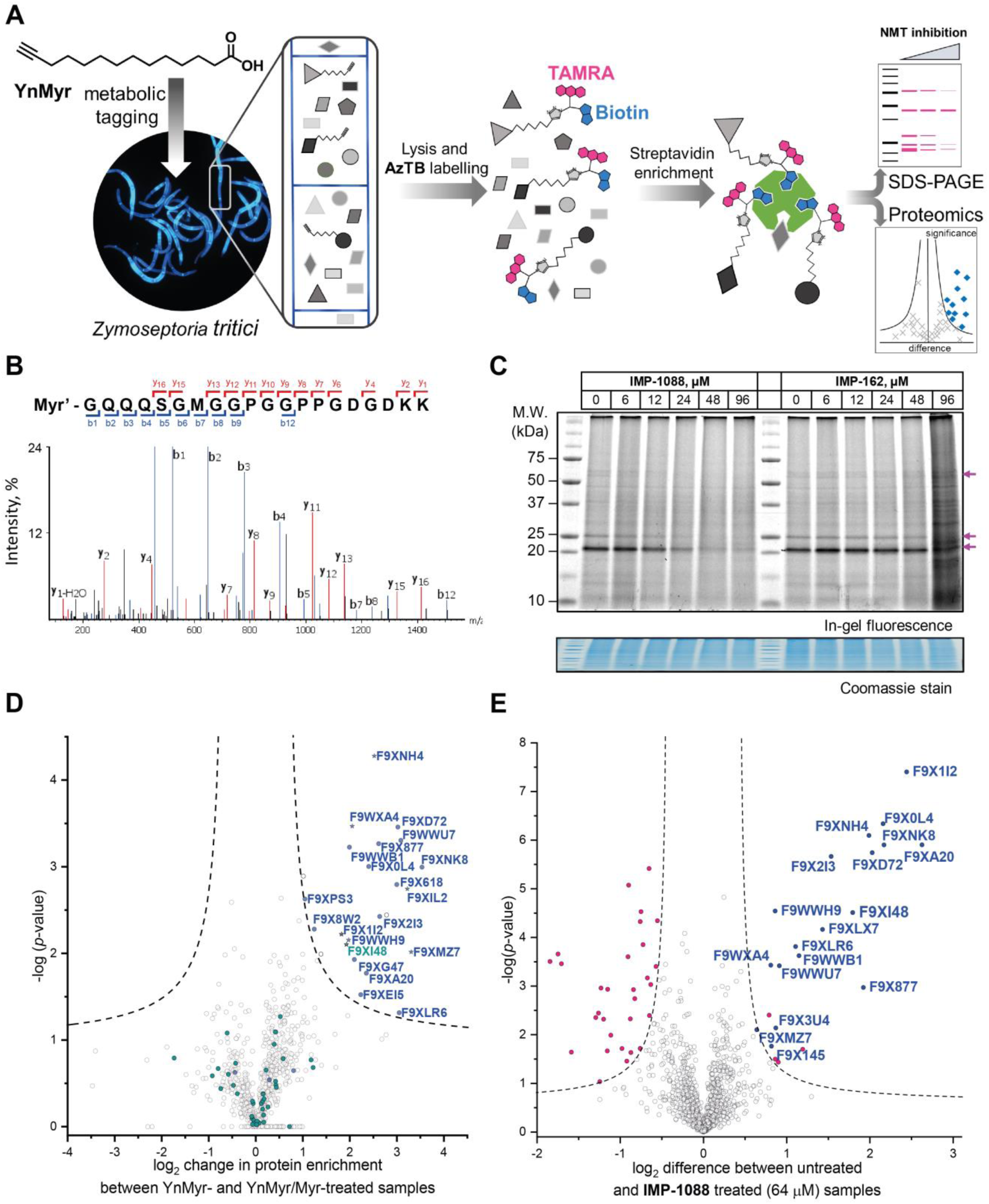
Inhibition and assessment of myristoylation *in vivo*. **A**) Generalized workflow for identification and quantification of myristoylated proteins. Cells were treated with YnMyr in culture, and cell lysates ligated to multifunctional reagents bearing a TAMRA dye for fluorescent in-gel imaging and/or biotin for affinity enrichment and analysis by proteomics. **B**) Fragmentation (MS/MS) spectrum of the N-terminal peptide of putative ZtNMT substrate F9XI48 showing fragmentation consistent with tagging by the YnMyr probe ligated to capture reagent (Myr’). **C**) Impact on myristoylation labeling by **IMP-1088** or **IMP-162** followed by TAMRA fluorescence; cells were treated with compound and YnMyr for 4 h, followed by lysis and CuAAC labeling with AzTB. Protein loading is normalized across samples (Coomassie blue stain), magenta arrows indicate putative myristoylated proteins. **IMP-1088** dose-dependently reduces labeling, whereas treatment with **IMP-162** does not; **IMP-1088** has no detectable effect on cell viability over 4 h, however the highest doses of **IMP-162** treatment result in cell death within the timeframe of the experiment, indicating an off-target mode of action for this compound. **D**) Identification of YnMyr-enriched ZtNMT substrates in *Z. tritici* by label-free quantification proteomics; proteins possessing an N-terminal glycine are shown in blue for proteins predicted to be myristoylated by the Myristoylator algorithm^34^, or green for those predicted to be non-myristoylated, myristoylated peptide was identified for the proteins marked as a star; dashed line represents permutation-based false discovery rate (FDR) of 0.05 (two-tailed *t*-test, 250 randomizations). **E**) Inhibition of ZtNMT substrates in *Z. tritici* analyzed by TMT proteomics; proteins identified as myristoylated (Table 1) are shown in blue, significant non-myristoylated proteins are indicated in magenta; dashed line represents permutation-based false discovery rate (FDR) of 0.05 (two-tailed *t*-test, 250 randomizations).

### Chemical proteomics provides the first insights into the *N*-myristoylated proteome in *Z. tritici*

In order to assess the potential of NMT as a target in *Z. tritici* we first aimed to characterize the *N*-myristoylation profile of *Z. tritici* in a native context. An alkyne-bearing analogue of myristic acid, YnMyr^28^, was used as described previously in protozoan parasites^29^. *Z. tritici* cultures were treated with YnMyr to label myristoylated proteins, with myristic acid added to compete for labeling in a negative control. Lysates treated with the YnMyr probe were subjected to Cu-catalyzed alkyne-azide cycloaddition (CuAAC) with capture reagent AzTB^30^ bearing both TAMRA fluorophore and biotin, or AzRB carrying biotin and an arginine cleavage site for enhanced identification of modified peptides. To specifically enrich labeled proteins, neutravidin-based magnetic beads were used for gel-based experiments and streptavidin-agarose resin was used in proteomics workflows^20,31^ (Figure 2A). SDS-PAGE separation and gel imaging showed that YnMyr produces concentration-dependent labeling that was outcompeted in the presence of myristic acid, the natural substrate of NMT (Figure S3A).

**Table 1.**
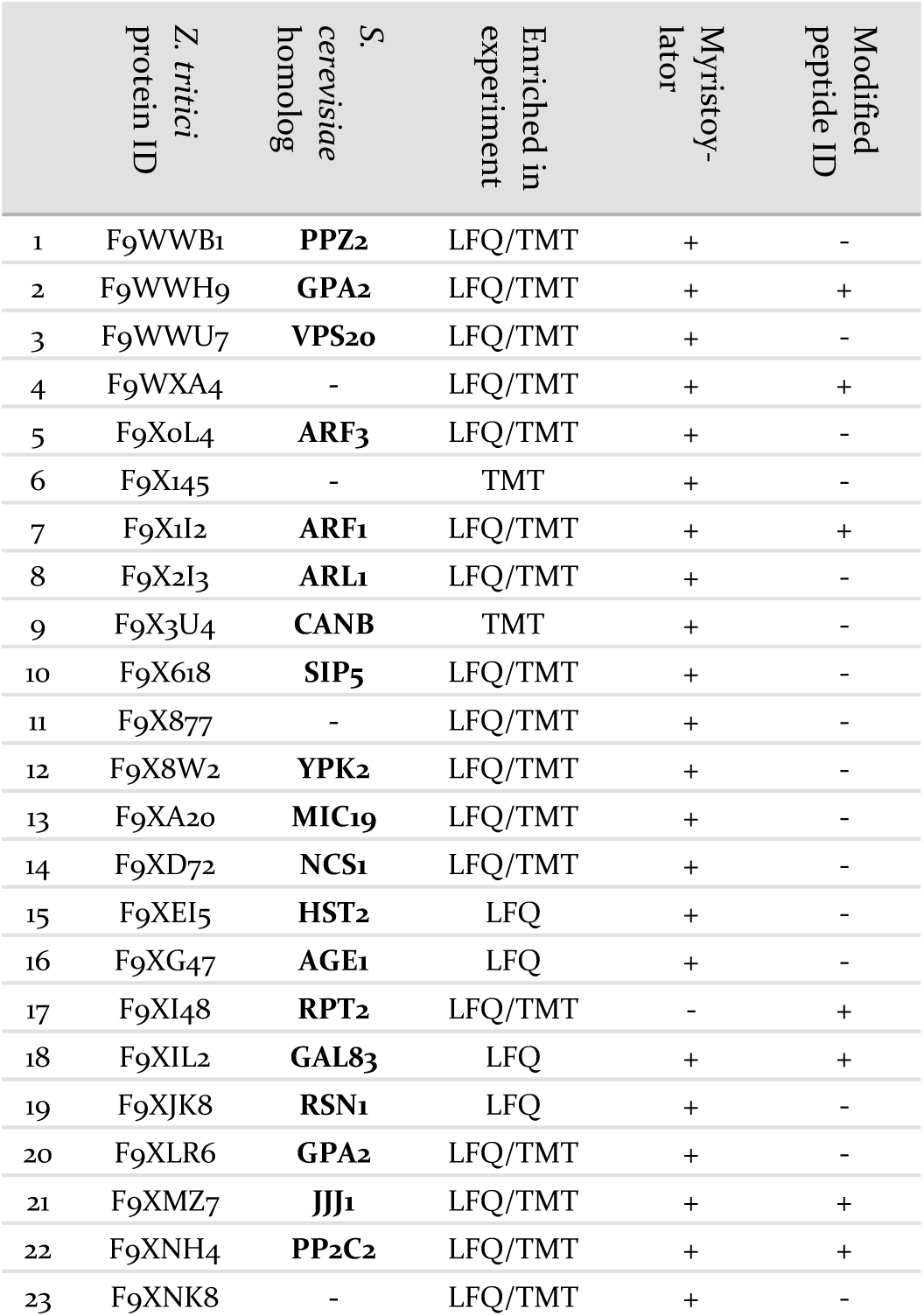
Newly identified *N*-myristoylated proteins in *Z. tritici*. UniProtKB primary accession numbers are used to name *Z. tritici* proteins. In the table are presented proteins with at least two independent lines of evidence of their myristoylation from either quantitative proteomics (indicated by the type of experiment with the positive identification, LFQ or TMT), *N*-myristoylated peptide ID or *in silico* prediction by Myristoylator^34^. *S. cerevisiae* homologs found by the whole-proteome BLAST+ software^39^ search are indicated.

From label-free mass spectrometry (MS) data, we identified multiple instances of N-terminal tryptic peptide from seven *Z. tritici* proteins modified with YnMyr and AzRB (Figures 2B, S3B and S3C) in MS/MS spectra, per-mitting direct assignment of these proteins as N-terminally myristoylated. For example, although *Z. tritici* protein F9XI48 was not predicted to be myristoylated based on the Uniprot sequence, we identified numerous high-quality spectra for F9XI48 N-terminal peptides modified with YnMyr-AzRB (Figure 2B). Note that UniProtKB primary accession numbers have been used to name *Z. tritici* proteins identified for the first time in these experiments. F9XI48 is a homolog of *S. cerevisiae* RPT2 and human PSMC1 proteins, which are myristoylated AAA-ATPases in the 26S proteasome complex, lending support to its assignment as a myristoylated protein in *Z. tritici*.

By quantitative analysis, we found 25 proteins significantly enriched relative to samples without myristic acid competition (Figure 2D). We excluded those lacking an N-terminal glycine which is known to be essential for NMT catalysis of *N*-myristoylation^35,36^, leaving 22 putative NMT substrate proteins; we speculate that non-myristoylated proteins may be enriched by non-covalent interactions, in line with observations in previous studies in human cells^31^. To complement the analysis, we applied a predictive web-based tool ‘Myristoylator’^34^ to analyze myristoylation by an orthogonal method; this predictor is trained primarily on an experimental yeast (*S. cerevisiae*) NMT substrate set, so we expected it to be a reasonable predictor of myristoylation for *Z. tritici*. Overall, out of all 794 proteins quantified in our proteomics analysis, N-terminal glycine was present in 66 proteins, and 25 were predicted to be myristoylated, 20 of which were enriched in the proteomics analysis (enrichment factor 26.5, −logP=30.8) indicating a high degree of fidelity between proteomic and *in silico* analyses. Annotation of the *Z. tritici* proteome remains under development in the current literature, as evidenced by the annotation of one identified myristoylated protein, F9XJK8, as a fragment in the Uniprot *Z. tritici* proteome^37^. However, a more recent annotation assigns a complete sequence to this genetic feature which contains an N-terminal glycine and is predicted myristoylated^38^, providing complementary lines of experimental and genomic evidence for both sequence and myristoylation state.

Finally, by analyzing YnMyr + DMSO vs YnMyr + Myr data quantified as part of a tandem mass tagging (TMT) experiment under the same conditions (see below), we found most of the myristoylated proteins already assigned in the label-free quantification (LFQ) experiment, as well as two additional proteins (Figure S4). Table 1 thus assembles the first experimentally determined *Z. tritici N*-myristoylated proteome, deduced from two independently replicated quantitative proteomics experiments, computational predictions, and direct MS/MS evidence.

### IMP-1088 inhibits N-myristoylation in the wheat pathogen *Z. tritici*

We next explored the mode-of-action for two potent *in vivo* compounds, **IMP-1088** and **IMP-162**. YnMyr was used to measure the effect of inhibitor on NMT activity in live *Z. tritici*. **IMP-1088** has no effect on cell viability over the short timeframe of the experiment (4 h), a phenotype consistent with the well-established dependence of *N*-myristoylation on de novo protein synthesis; cytotoxicity typically results from progressive reduction in *N*-myristoylation over time^21,31^. In contrast, treatment with **IMP-162** did not produce any reduction of in-gel band intensities (Figure 2C) but showed a remarkable ability to kill *Z. tritici* culture just after four hours of treatment in logarithmic growth phase. At concentrations above EC50 (24 µM and 48 µM) parts of the culture produced a phenotype consistent with cell death as observed under a microscope, while at 96 µM most cells appear dead (data not shown). Combined with the low activity against ZtNMT, this divergent phenotype further confirms an off-target mode of action which may merit future investigation, given the *in vivo* potency of this compound.

In contrast, **IMP-1088** reduced YnMyr tagging (Figure 2C) with half maximal concentration (TC50^20^) of 14 ± 1 µM measured by quantifying intensity of the most prominent band (ca. 20 kDa) by in-gel fluorescence, which correlates well with the **IMP-1088** antiproliferative EC50 value of 23 μM. To investigate the effect of **IMP-1088** on the myristoylated proteome we treated *Z. tritici* cultures with YnMyr and increasing concentrations of **IMP-1088** and processed samples for proteomic analysis as described above. Tryptic peptides were labelled with 6-plex TMT reagents; all six samples were treated with YnMyr, and additionally with DMSO (vehicle), or 8, 16, 32, or 64 µM **IMP-1088**, or with myristic acid. Myristic acid was used as a point of reference for the previous LFQ analysis and as a negative control treatment which competes with alkyne-bearing YnMyr for myristoylated proteome labeling. Quantitative differences between 64 µM **IMP-1088** and DMSO-treated samples strongly indicated an NMT-dependent mode-of-action, with most of the myristoylated proteins showing reduced tagging with YnMyr alkyne in the presence of NMT inhibitor (Figure 2E).

Analyzed by unbiased hierarchical clustering, inhibitor-treated samples with increasing concentrations of **IMP-1088** grouped naturally into a single distinct cluster (Figure S5 Cluster 1). Within Cluster 1 we identified three subsets of myristoylated proteins that respond differentially to NMT inhibition: with TC50 (concentration of half-tagging with YnMyr) in the ranges 8-16 µM, 16-32 µM and 32-64 µM, respectively, for each fraction. A range of differential responses has been observed previously in human and parasite cells, and likely results from differential efficiency of myristoylation between substrates under inhibition^29,31,40^.

Other proteins with statistically significant trends identified by analysis of variance (ANOVA) test were grouped into Clusters 2, 3 and 4 (Figure S5). Proteins in these clusters showed ether increased or decreased intensities of signals in inhibitor-treated samples but were unchanged in the negative control (myristic acid-treated samples), indicating that these proteins are non-specific binders, and that changes in enrichment are unrelated to the labeling of *N*-myristoylated proteins. In the same fashion, non-myristoylated proteins with significant change in abundance between samples treated with DMSO and 64 µM **IMP-1088** are indicated in magenta on Figure 2E. We hypothesized that changes in abundances of these proteins are not due to differential enrichment, but due to either up- or down-regulation under inhibitor treatment; such large changes over a 4 h timeframe are in line with the fast division time of *Z. tritici* (5-6 hours), and much faster than those seen in organisms with a slower cell cycle^41,42^. These data prompted us to investigate inhibitor-treated samples omitting the enrichment step and analyzing proteome-wide changes.

### IMP-1088 acts on ZtNMT to induce a potent stress response *in vivo*

Whole proteome analysis of *Z. tritici* cultures revealed significant dose-dependent changes in the expression of multiple proteins in response to **IMP-1088** (Figure 3A, Table S2). Limited annotation of *Z. tritici* genes^43^ prompted us to undertake an in-depth analysis using NCBI BLAST+^39^ to find closest homologs in *S. cerevisiae*, one of the most well-annotated yeast genomes available. We performed the search filtering out poor alignment matches (e-value > 0.01) and selected a single best match in the *S. cerevisiae* proteome for each *Z. tritici* protein. Approximately 56% (6168 out of 10972 proteins) of *Z. tritici* proteins were found to have reasonable alignment to a *S. cerevisiae* protein and were assigned Kyoto Encyclopedia of Genes and Genomes (KEGG) terms and Gene Ontology (GO) terms based on the relevant *S. cerevisiae* terms (Table S3).

**Figure 3.**
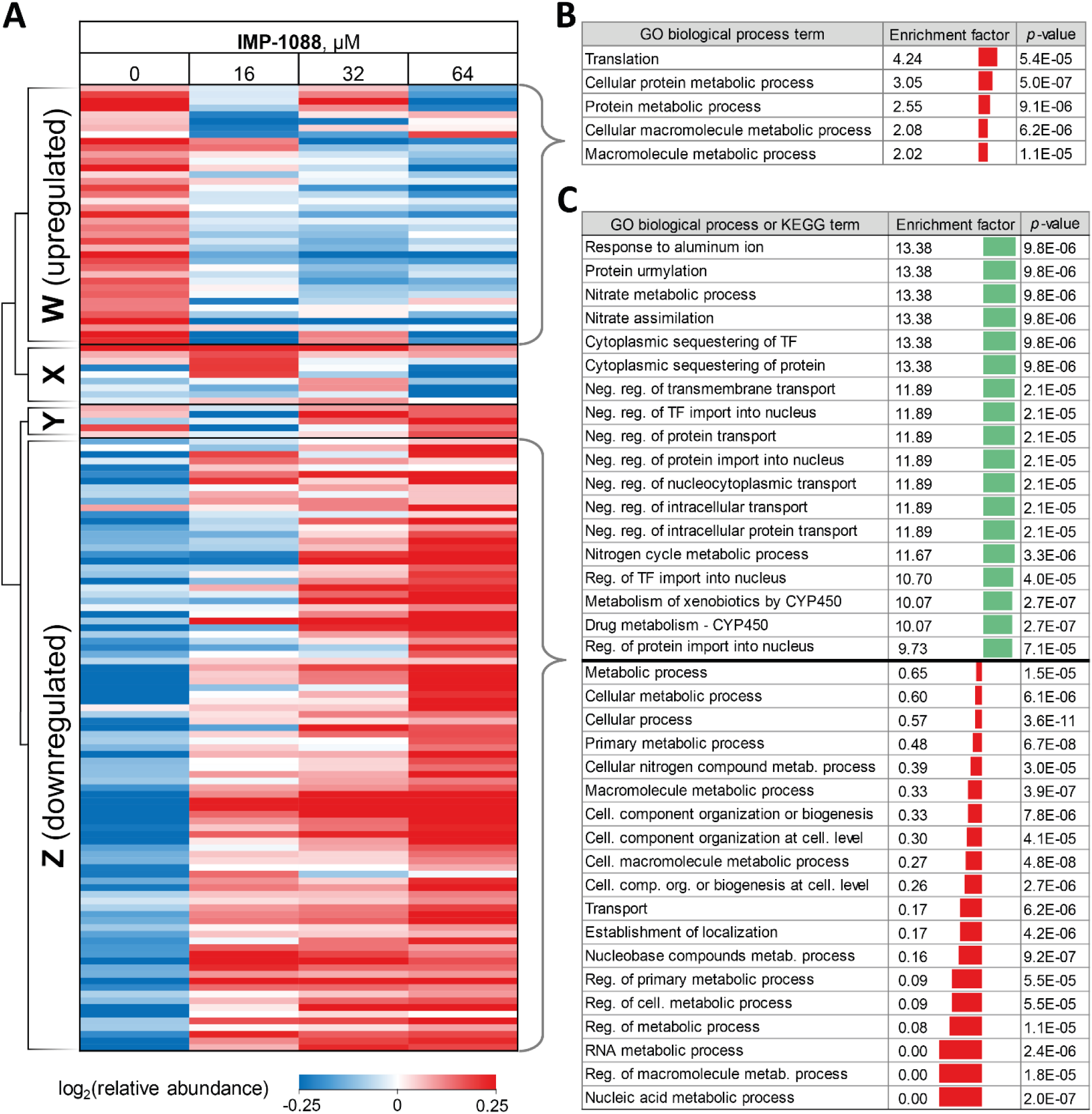
Response of *Z. tritici* to treatment with NMT inhibitors. **A**) Heatmap showing proteins that exhibit significant changes (ANOVA test FDR=0.01) in expression levels in the whole lysate in response to increasing **IMP-1088** concentrations over 4 h. Proteins downregulated upon **IMP-1088** treatment are grouped by hierarchical clustering by Pearson correlation into Cluster W. Upregulated proteins group into Cluster Z. Inconsistently changing abundance proteins are in Clusters X and Y. Higher protein abundance (red) to lower protein abundance (blue) values are indicated by color scale at the bottom. Data is normalized by subtraction of the row mean values. **B**) GO biological process and KEGG term enrichments using Fisher’s exact test applied to proteins in Cluster W. **C**) GO biological process term enrichments from Fisher’s exact test applied to proteins in Cluster Z. GO annotation were assigned to *Z. tritici* proteins based on *S. cerevisiae* homolog (BLAST analysis) terms. Enrichment factor indicator bar is colored red for GO terms over-represented among down-regulated and under-represented among up-regulated proteins; the indicator bar is green for GO terms over-represented among up-regulated proteins.

For 39 proteins in Cluster W (Figure 3A, proteins down-regulated upon inhibitor treatment), 5 Gene Ontology Biological Process (GOBP) terms were found to be over-represented (FDR < 0.02, Figure 3B) across the entire range of **IMP-1088** concentrations. Four of these terms are redundant and are processes associated with metabolism of macromolecules or proteins and one term associated with translation. Similarly, for 92 proteins in Cluster Z (Figure 3A, proteins upregulated upon inhibitor treatment) we found 35 GOBP terms either over- or under-represented (FDR < 0.02, Figure 3C). Terms under-represented in Cluster Z proteins mirror those over-represented in Cluster W: transport, biogenesis and metabolic processes are suppressed indicating *Z. tritici* may be indeed switching from the logarithmic expansion to a dormant phase. These data indicate a widespread decrease in metabolic and proliferative activity upon **IMP-1088** treatment.

We also found several GO terms to be over-represented using Fisher’s exact test, based on proteins in Cluster Z. Processes related to sequestering (GO:0051220, GO:0042994), nitrogen metabolism (GO:0042126, GO:0042128, GO:0071941) negative regulation of protein transport (GO:0034763, GO:0051224, GO:0032387, GO:0042308, GO:0042992, GO:0046823, GO:0090317) were found to be the most significantly up-regulated (Figure 3C). We noticed a significant overlap in the annotation terms indicated in Table 2. In every category five proteins remained the same (namely, F9WWN5, F9×5T1, F9XF90, F9XGG1 and F9XLJ2) comprising these categories together with additional proteins. This overlap stems from the annotation of these proteins derived from homology in *S. cerevisiae*. All five proteins, and also three additional non-significantly enriched proteins in the proteomics analysis are analogues of *S. cerevisiae* URE2, and thus inherit URE2 annotations. Such extensive homology is not surprising given identification of expanded URE2-analogue families in various saprophytic and wood-decaying fungi, which have been linked to metabolism of diverse substrates by these fungi^44–46^. Furthermore, a BLAST homology search found two additional URE analogues (10 in total) among proteins which were not identified by our proteomics analyses. These proteins share a high degree of similarity to the *S. cerevisiae* URE2 (Table S3).

Among URE2 functions reported to date are control of transcription factor GLN3 in response to available nitrogen sources^47^, peroxidase activity^48^, heavy metal detoxification^49^, self-propagating prion protein upon conformational changes^50^ and small-molecule binding through its GST domain^44,51^. Among the eight proteins enriched that are KEGG cytochrome P450-related (metabolism of xenobiotics and drugs metabolism), we additionally identified analogues of *S. cerevisiae* glutathione *S*-transferase 1 (GTT1), and alcohol dehydrogenases 2 and 5 (ADH2 and ADH5).

Taken together, these data support the hypothesis that in response to NMT inhibitor *Z. tritici* rapidly, within four hours of treatment, over-expresses numerous small molecule metabolizing enzymes, among them cytochrome P450s and glutathione *S*-transferases, to combat the negative impacts of the chemical inhibitor. We hypothesize that this may lie at the origin of the >1000-fold decrease in the inhibitor activity translation from ZtNMT enzyme inhibition to *Z. tritici* culture inhibition.

## DISCUSSION

Agricultural pathogens are under-represented in the current literature relative to their impact on society and the requirement for advances in crop protection. Our study demonstrates the power of chemical proteomics for understanding drug mode of action in the causative agent of a highly damaging wheat disease, Septoria tritici blotch. Extending three previous studies that have focused on *Z. tritici* secretome^43^, phosphoproteomics^52^, and whole proteome analysis^53^, our functional proteomics study in *Z. tritici* demonstrates the applicability of chemical biology and medicinal chemistry approaches to understanding this important crop pathogen. We also demonstrate that the lack of basic functional genome annotation in this organism can be overcome by comparative bioinformatics approaches. The well-annotated *S. cerevisiae* genome has previously been used to re-annotate closely-related fungal organisms^54^, and here we extend this functional annotation approach to *Z. tritici*, a more distant *S. cerevisiae* relative that shares only the common phylum of Ascomycota, uncovering numerous biological processes underlying *Z. tritici* response to a chemical inhibitor. The powerful response elicited by *Z. tritici* to a novel foreign compound may also render it a valuable biological model to study metabolism and stress response to chemical inhibitors on the systems level.

Compounds identified in this study include several of the most potent fungal NMT inhibitors reported to date and rationalized their binding mode and SAR using dynamic molecular models. Through chemical proteomic approaches we investigated the origin of the prominent drop in potency on moving from enzyme IC_50_ to whole-cell culture EC_50_ values. It appears that inhibitors from existing NMT inhibitor development pipelines using generic parameters for physicochemical optimization are not ideally suited to the requirements of *Z. tritici* whole-cell activity. Based on our functional proteomic analyses, we hypothesize that it will be important for fungal NMT inhibitors to withstand aggressive *Z. tritici* metabolic and cellular efflux mechanisms. As part of the screening we serendipitously discovered a potent whole-cell fungal inhibitor **IMP-162** of unknown mode-of-action, capable of killing *Z. tritici* cultures growing in logarithmic phase in rich media within 4 h of treatment. A further investigation of activity of **IMP-162** is beyond the scope of the present study, but merits future investigation.

We have shown that NMT inhibitors restrict *Z. tritici* proliferation in the micromolar concentration range. We demonstrate that the most potent of these inhibitors acts on-target, and further investigated the mechanisms of *Z. tritici* response to inhibitor, describing a defense mechanism employed to withstand inhibition. Our evidence suggests that this putative defense mechanism deviates from conventional understanding that metabolism primarily occurs via cytochrome P450 mediated oxidation. Instead, it appears that *Z. tritici* upregulates expression of a family of URE2-related glutathione *S*-transferases, which have been previously implicated in the metabolism of small molecules. Proliferation and metabolism are rapidly decreased in *Z. tritici* upon treatment with NMT inhibitor, a response which may present challenges in targeting this agricultural pathogen with modes of action which are most effective on actively growing cultures.

Large decreases of activity have been reported for other NMT inhibitors previously. For example, a benzothiazole-based *Candida albicans* inhibitor was found to be >1500-fold less active at inhibiting fungal growth compared to its enzymatic activity against CaNMT^55^, whilst a pyrazole sulfonamide-based *Aspergillus fumigatus* NMT inhibitor similar to **IMP-366** suffers a >80000-fold drop in activity^17^. Interestingly, *Ure2* in the related fungus *Aspergillus nidulans* was shown to a be the key gene involved in defense against a variety of toxic small molecules^56^. This indiscriminate defense mechanism appears unlikely to be mediated by NMT, which points to the hypothesis that **IMP-1088** upregulates URE2 and its analogues through an alternative interaction. The full catalogue of enzymes participating in the degradation of targeted inhibitors in *Z. tritici* remains to be established, and the currently very limited annotation of the *Z. tritici* genome significantly restricts further analysis. It could be anticipated that inhibition of the most important metabolizing enzymes and pathways could increase sensitivity of *Z. tritici* to antifungals, which may warrant further investigation of GST and CYP450 co-inhibition cooperativity. The small-molecule metabolism properties of GSTs have so far received much less attention compared to CYP450, and this works provides an incentive to investigate their role in future work.

## METHODS

### Recombinant ZtNMTΔ99 preparation

ZtNMTΔ99 in pET24a vector was transformed into BL21 (DE3) *E. coli* cells, for shake flask expression. Cells were grown at 37°C 200 rpm in auto-induction media to an OD600 of approx. 0.6 before lowering the temperature to 20°C for overnight expression. Harvested biomass was resuspended in 5 volumes of lysis buffer containing 100mM Tris pH 7.5, 200 mM NaCl, 20 mM imidazole plus protease inhibitors. Resuspended cells were lysed using the Constant Systems cell disruptor at 20000 psi (two passes). The cell lysate was clarified by centrifugation and the supernatant was applied to a GE 5 ml HisTrap FF column equilibrated in IMAC buffer A 100 mM Tris pH 7.5, 200 mM NaCl, 20 mM imidazole. The column was washed with 20 CVs of equilibration buffer. Bound protein was eluted using IMAC Buffer B 100 mM Tris pH 7.5, 200 mM NaCl, 300 mM imidazole over a 3.5 CV step gradient. Eluted protein was loaded onto a GE 26/60 Superdex 200 SEC column equilibrated in 20 mM Tris pH 7.5, 100 mM NaCl. Pooled fractions were concentrated in a 10 kDa cut off Vivaspin and 10% glycerol added. Protein concentration was determined using the Nanodrop ME52070. The final sample was run on an SDS-PAGE gel and purity determined using the Bio-Rad ChemiDoc MP Imaging system with ImageLab software. Purity was found to be approximately 99%.

### ZtNMT IC_50_ assay

The assay was adapted from a previous publication^25^. Briefly, a solution of an inhibitor (variable concentration), myristoyl-CoA (4 μM), ZtNMTΔ99 (200 ng ml^-1^), peptide substrate (ZtARF2 N-terminal sequence: NH_2_-GLTFSKLF-OH, 8 μM), 7-diethylamino-3-(4′-maleimidylphenyl)-4-methylcoumarin (CPM, 8 μM) dye were mixed on a 96-well plate, each condition in duplicate. Accumulation of fluorescence signal at 470 nm was measured at one-minute intervals during the first 20 minutes of the reaction. Rates of the enzymatic reaction were calculated from the linear range of the resulting fluorescence over time curve using GraphPad Prizm 5. Then, sigmoidal dose-response (variable curve) fitting was applied to the data to obtain IC_50_ values. The assay was performed with at least 2 replicates, mean and confidence intervals are reported.

### Anti-proliferative EC_50_ assay

Promega’s CellTiter 96® Aqueous Non-Radioactive Cell Proliferation Assay was used following the instruction manual. Briefly, 1*10^7^ ml^-1^ of *Z. tritici* spores were seeded on 96-well plate in 100 μl Vogel’s Minimal Media (VMM) with a corresponding concentration of an inhibitor and kept in a shaking incubator at 24°C at 140 RPM for 40 h. At least three replicates per each condition were set up. 20 μl of PMS/MTS solution was added to each well and the plate was left stirring for further 3 h. Then, absorbance at 490nm was measured and the resulting values were fitted using sigmoidal dose-response (variable curve) in GraphPad Prizm 5 to produce EC_50_ values. The assay was performed with at least 4 replicates, mean and confidence intervals are reported.

### Click chemistry

Click chemistry was performed as described in Thinon et al. 2014^31^. Briefly, 0.06 volumes of click master mix was prepared by combining 0.01 volume of 10 mM in DMSO of the azide reagent of choice, 0.02 volumes of 50 mM copper sulfate in MilliQ water, 0.02 volumes of 50 mM TCEP in MilliQ water and 0.01 volume 10 mM TBTA in DMSO. The amount of lysate corresponding to a concentration of 2 mg ml^-1^ of protein in 1 volume of final mixture was taken and diluted with lysis buffer to 0.94 volumes. 0.06 volumes of click master mix was added to 0.94 volumes of protein solution and the mixture was stirred for 1 h in bench vortex. After, the reaction was quenched by addition of 0.02 volumes of 0.5 M EDTA. 2 volumes of methanol followed by 0.5 volumes of chloroform and followed by 1 volume of MilliQ water were then added to precipitate proteins. The precipitate was pelleted by centrifugation at 40 °C for 10 min at 17000 g. The pellet was then washed twice with cold methanol by resuspension and centrifugation. Finally, the pellet was dissolved in 0.1 volume of 2% SDS/PBS and diluted by 0.9 volumes of PBS.

The click reagents used are described in Broncel et al. 2015. We used AzT for in gel fluorescence analysis, AzTB for enrichment on the streptavidin beads and following in gel fluorescence analysis or western blotting, AzRB for enrichment on neutravidin beads and proteomics analysis.

### Streptavidin enrichment

15 μl of Dynabeads™ MyOne™ Streptavidin C1 from Invitrogen (streptavidin beads) were washed three times with 100 μl 0.2% SDS in PBS. Then 100 μl of 1 mg ml^-1^ solution of proteins after click reaction with AzTB was added and the suspension was stirred for 2 hours at room temperature. The beads were washed three times with 1% SDS in PBS and then resuspended in 15 μl of 1x SDS PAGE Sample Loading Buffer. The tubes were placed in a heating block at 950C for 10 minutes. The supernatant was then loaded onto 12% polyacrylamide gel.

### SDS-PAGE and in-gel fluorescence

Components were purchased from National Diagnostics and the gels were cast according to manufacturer’s protocol. The gels were run for 15 min at 80 V followed by 60 min at 160 V in 0.192 M glycine, 25 mM Tris base, 1% SDS buffer. Fluorescent images of the gels were recorded on a GE Typhoon FLA 9500 equipped with 532 nm excitation laser and an LPG emission filter. After the imaging, the gels were placed in a Coomassie staining buffer (10% w/v ammonium sulfate, 10% v/v phosphoric acid, 20% v/v methanol, 1.2% w/v Coomassie brilliant blue G) overnight. The gels were then washed in MilliQ water for 1 h and images were scanned using a Canon LiDE 120 office scanner.

### Prediction of the myristoylation sites

Myristoylator is a computational tool publicly available at http://web.expasy.org/myristoylator/ (Bologna et al. 2004).

### Homology mapping of *Z. tritici* proteins to *S. cerevisiae* proteins

NCBI BLAST^39^ 2.6.0+ was used to perform homology search with the following settings: the maximum e-value of a hit should not exceed 0.01 and only one top hit was retained. Results of the search are available in the Table S3, where each match is annotated with the percentage and the number of positions aligned, e-value and score.

### Sequence alignment and phylogenetic tree

Sequence alignment and phylogenetic tree were created using CLC sequence viewer 7.5 default settings. The protein sequences used were downloaded from uniprot.org.

### ZtNMT homology model

A ZtNMT homology model was built using the SWISS-Model online suite. The model was created based on the *A. fumigatus* NMT X-Ray structure 4UWI (Brand et al. 2014) using the ZtNMT^132-535^ sequence and using the default SWISS-Model settings.

#### ZtNMT^132-535^

EGPILPPTVCKKVAKPELEKLVDGFEWCGIDLEDKEELQEFYDLLYNHYVEDTEGSFRFNYSKEFLAWALKPPGWTKECHIGVRTKTGDDGKKGKLVASIAGIPVSLTVRGKQVDACEINFLAIHRKLRNKRLAPVLIKEVTRRYYLNGIYQALYTAGTLLPTPVSTCRYYHRSLDWEHLYKNGFSHLPPHSSELRMKLKYKLEDKTALKGLRPMKPADIPAVKELLSRYNERFHLRQNFTEEDVAHYLCSDISKGVVWSYVVEEKGKITDFISYYLLESTVLKSSNKRETIRAAYLYYYASDSAFPSSTSKAPSNSTQNALQARLQLLVHDALILAKKDDFHVFNALTLLDNPLFLKEQKFEPGDGKLHYYLFNWRTESLNGGVDERNQIDVTKMGGVGVVML.

### Proteomics

All methods related to proteomics sample preparation, mass spectrometry and data analysis are described in Supporting Information.

## ASSOCIATED CONTENT

### Supporting Information

This material is available free of charge via the Internet:

- Supporting Information (pdf) containing Figures S1-S5, Table S1 and additional methods.
- Table S2. Proteomics data (xlsx) containing complete proteomics data table
- Table S3. Z. tritici homology mapping to S. cerevisiae proteome (xlsx) with results of *Z. tritici* proteins homology mapping.

## AUTHOR INFORMATION

### Author Contributions

R.O.F planned and undertook experiments and analyzed data. C.A.O. performed computational modeling experiments. B.S., J.M. and A.J.C. cloned and purified *Z. tritici* NMT protein. E.W.T. conceived the project and directed the research. R.O.F. and E.W.T. wrote the manuscript, with input from all the authors.

## ACKNOWLEDGMENT

The authors thank Dr Andrew Bell for his critical reading of the manuscript.

## ABBREVIATIONS

ANOVA: analysis of variance;
BLAST: Basic Local Alignment Search Tool;
CoA: coenzyme A;
CPM: 7-diethylamino-3-(4’-maleimidylphenyl)-4-methylcoumarin;
CuAAC: Cu-catalyzed alkyne-azide cycloaddition;
DMI: demethylase inhibitor;
DMSO: dimethyl sulfoxide;
EC50: half-maximal effective concentration;
FDR: false discovery rate;
GO: Gene Ontology;
GOBP: Gene Ontology Biological Process;
GST: Glutathione S-transferase;
IC50: half-maximal inhibitory concentration;
KEGG: Kyoto Encyclopedia of Genes and Genomes;
LFQ: label-free quantification;
MBC: methyl benzimidazole carbamate;
MS: mass spectrometry;
MTS: 3-(4,5-dimethylthiazol-2-yl)-5-(3-carboxymethoxyphenyl)-2-(4-sulfophenyl)-2H-tetrazolium;
Af/Ca/Hs/ZtNMT: *N*-Myristoyl transferase of *A. fumigatus*/*C. albicans*/*H. sapiens*/*Z. tritici* species;
QoI: quinone outside inhibitor;
RMSD: root-mean-square deviation of atomic positions;
SDHI: succinate dehydrogenase inhibitor;
STB: *Septoria tritici* blotch;
TC50: half-maximal tagging concentration;
TMT: tandem mass tagging.

