## Supporting information for "Wheat pathogen *Zymoseptoria tritici N*-myristoyltransferase inhibitors: on-target antifungal activity and an unusual metabolic defense mechanism"

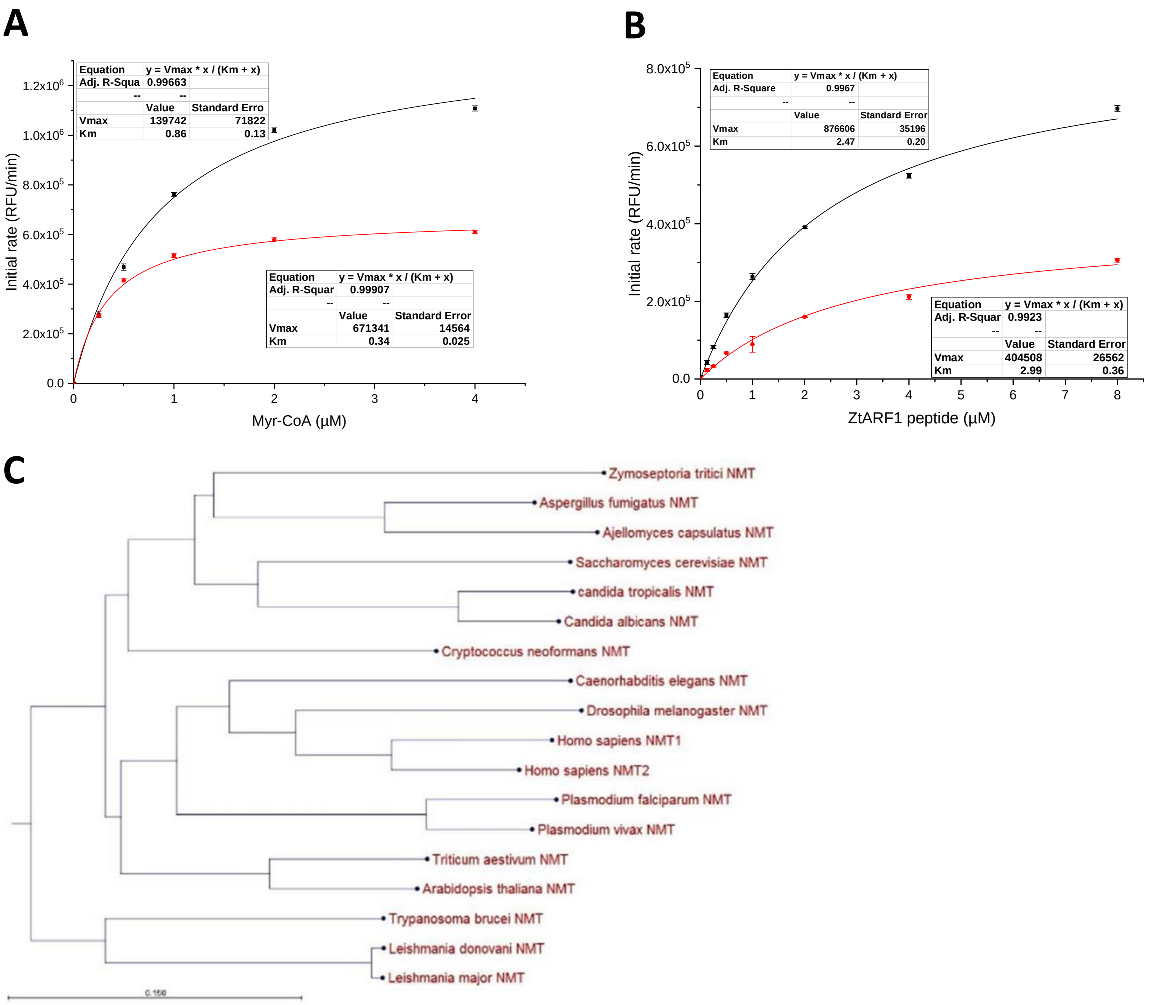

**Figure S1. ZtNMT characterization.** $K_M$ and $V_{max}$ constants measurements were performed at two ZtNMT concentrations: 100 ng/ml and 200 ng/ml. A). Myristoyl-CoA $K_m$ and $V_{max}$ measured at 8 µM (saturating concentration) of ZtARF1 $NH_2$-GLTFSKLF-OH peptide. B) ZtARF1 $K_m$ and $V_{max}$ measured at 4 µM (saturating concentration) of Myristoyl-CoA. The kinetic constants for both NMT substrates in ZtNMTΔ99 at saturating concentrations of the complementary substrate are: $K_M(Myr\text{-}CoA)$ = 0.60 µM in the presence of 16 µM peptide, and $K_M(peptide)$= 2.73 µM in the presence of 32 µM myristoyl-CoA. C). ZtNMT clusters together with other fungal NMTs that were extensively studied. The closest evolutionary related NMT within the species shown appears to be *A. fumigatus* NMT.

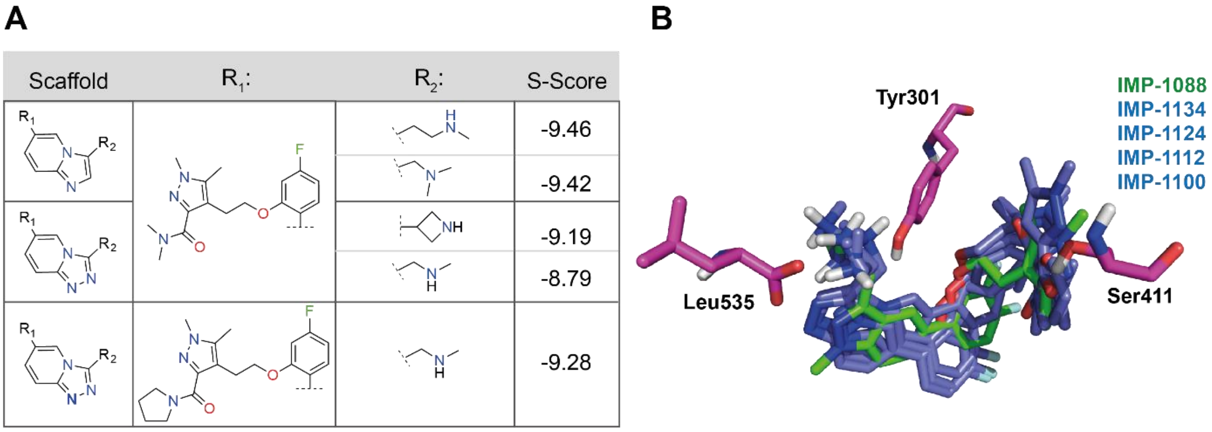

**Figure S2.** SAR analysis of MOE-docked IMP-derivatives. A). A MOE SAR report was generated by setting the S-score to activity (pKI/PIC50). The highest-ranking core scaffolds are listed along with their highest ranking $R_1$ and $R_2$ groups. Protonation state omitted. B). The docking poses of the top four hits (**IMP-1124**, **IMP-1134**, **IMP-1112** and **IMP-1100**) were overlaid with docked **IMP-1088** in the ZtNMT homology model.

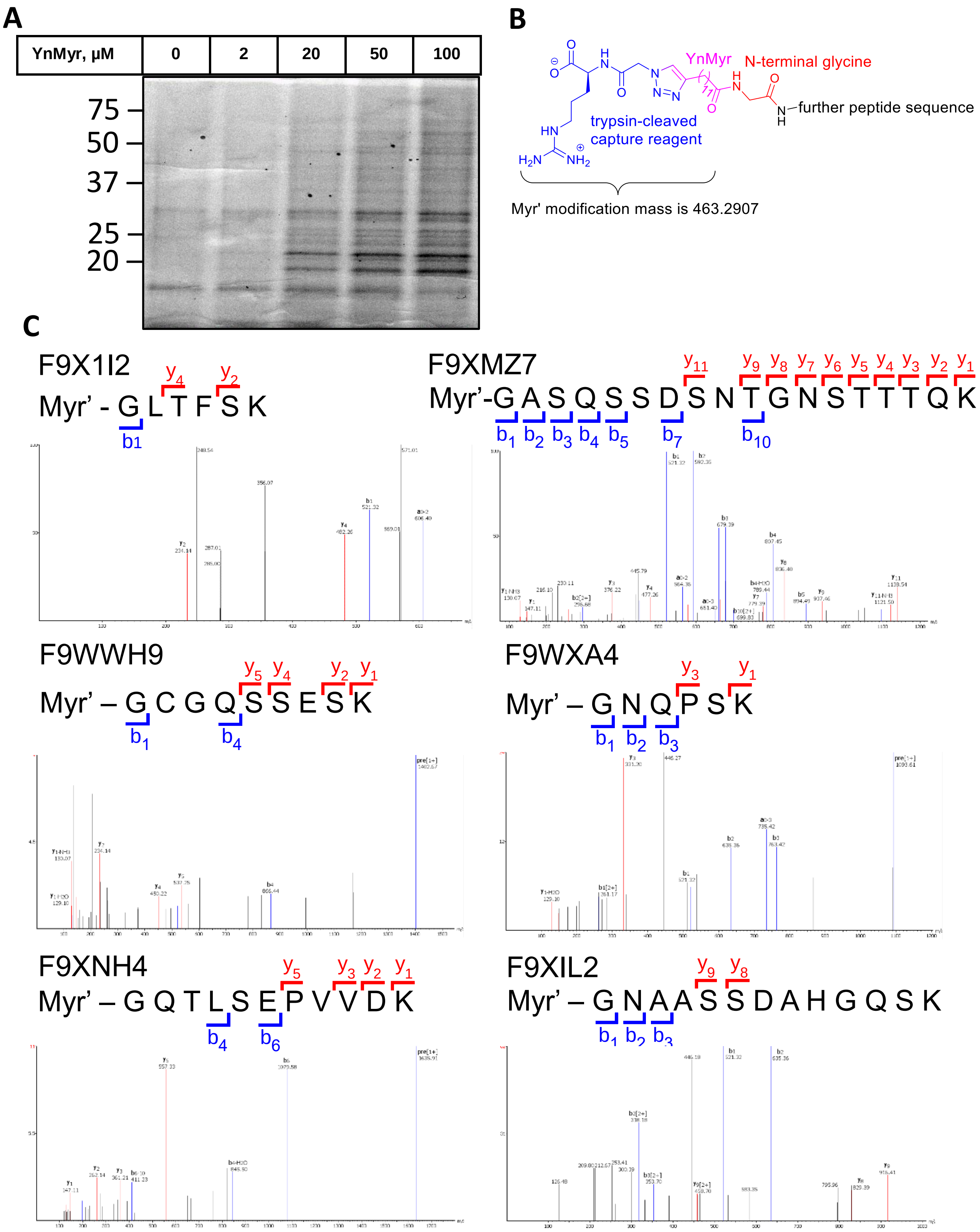

**Figure S3. MS/MS spectra of Myr'-modified (myristoylated) peptides identified by proteomics.** A). Concentration-dependent labelling with YnMyr/AzT in *Z. tritici* lysates. *Z. tritici* cultures were treated with YnMyr for 4 h, lysed labelled with AzT and visualized via in-gel fluorescence. B). In MS/MS spectra, the Myr' modification is introduced to myristoylated proteins instead of the native myristate via YnMyr and identified upon labelling with capture reagent AzRB, enrichment and tryptic digestion followed by proteomics. C). MS/MS spectra of the myristoylated peptides identified in the enriched label-free proteomics. The *S. cerevisiae* homologues found for these proteins are as follows: F9X1I2 – ARF1, F9XMZ7 – JJ1, F9WWH9 – GPA2, F9WXA4 – no match, F9XNH4 – PP2C2, F9XIL2 – GAL83.

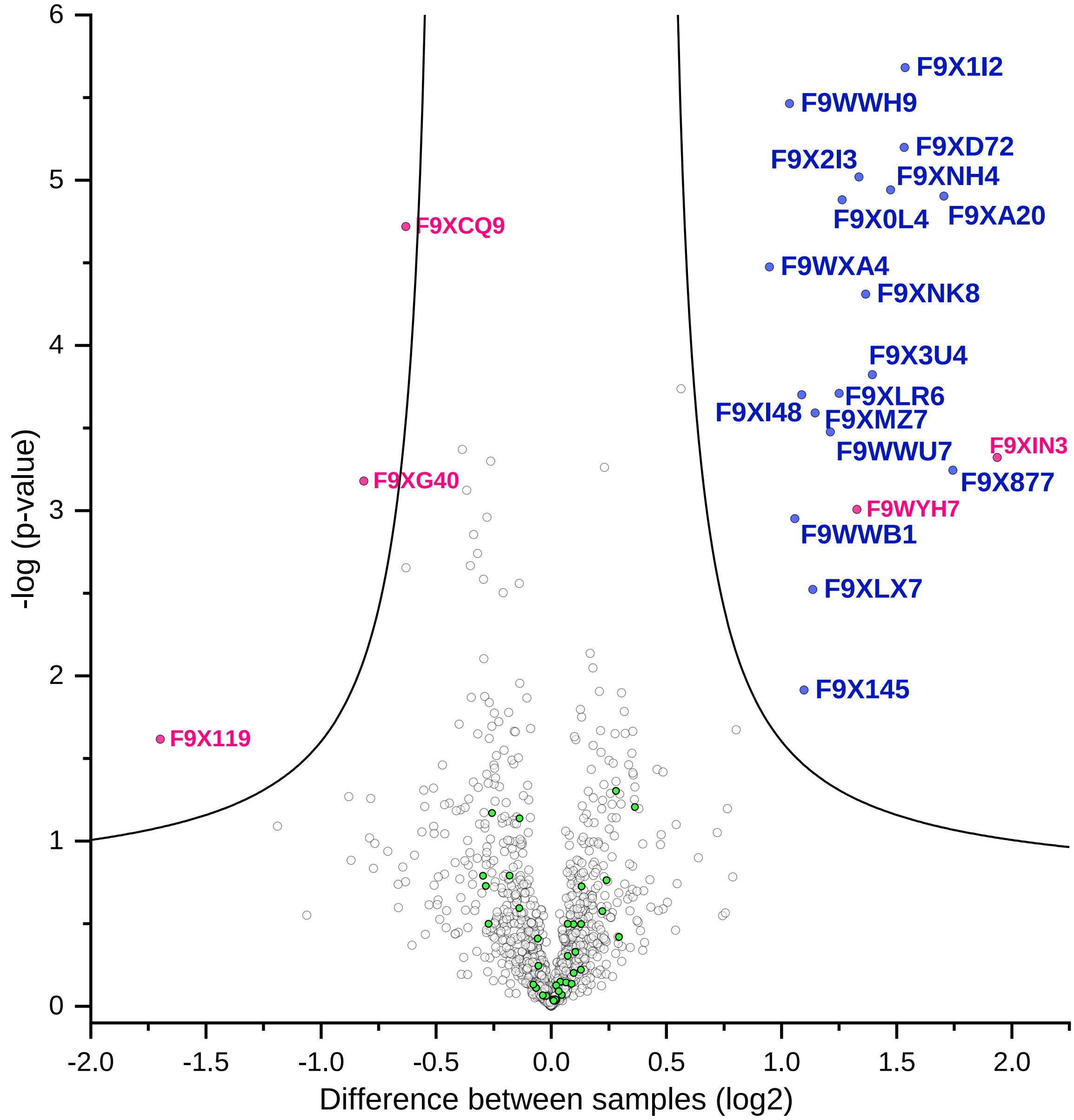

**Figure S4. Identification of additional myristoylated substrates in *Z. tritici* via the TMT proteomics.** In green are proteins with N-terminal glycine, in blue are the proteins predicted myristoylated by the "Myristoylator", in red are other proteins identified as significant.

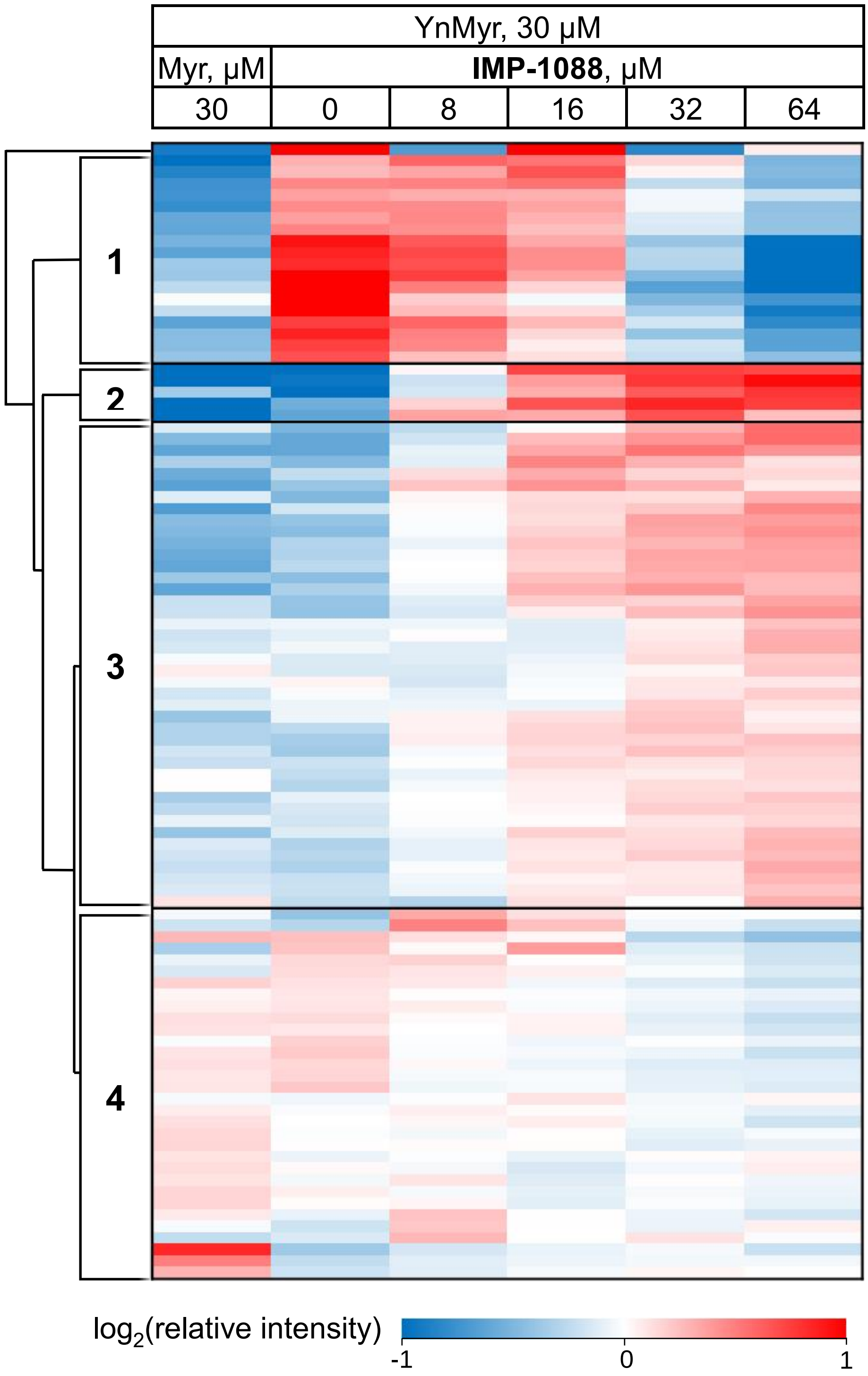

**Figure S5. Myristoylation enrichment proteomics upon NMT inhibition.** Heatmap shows proteins that exhibit significant changes (ANOVA test FDR=0.01) in enrichment upon treatment with YnMyr and different **IMP-1088** concentrations over 4 h. Myristoylated proteins are grouped in Cluster 1, They are abundant in YnMyr-treated samples, but their enrichment decreases upon competition with myristic acid (Myr) or NMT inhibition with **IMP-1088**. Higher protein abundance (red) to lower protein abundance (blue) values are indicated by color scale at the bottom. Data is normalized by subtraction of the row mean values.

### Supplemental methods

**Docking IMP-Derivatives in the ZtNMT Homology Model Binding-Pocket.** NMT inhibitors were selected based on structural diversity around three cored scaffolds, limited to compounds containing 1-*H*-indazole, triazolopyridine and imidazopyridine scaffolds. The compounds were rendered as 2D images using ChemDraw Professional (version 15.1.0.144) software, converted to SDF format, and then prepared for docking using Molecular Operating Environment (MOE 2018.01) software. After loading the SDF file, it was converted to a MOE Database file (MDB) and processed as follows: the compounds were washed, partial charges added (Amber10 forcefield) and energy minimized. Protonation states were enumerated and only non-cyclical nitrogens were protonated based on their protonation state between pH 7 – 8. To prepare the receptor protein, the ZtNMT PDB file, containing energy minimized IMP-1088, was loaded into MOE and processed using the "add hydrogens" and "connect and type" commands. Finally, the potential was adjusted using the "fix" command and the binding-site identified by selection of IMP-1088 ligand atoms already included in the ZtNMT homology model. The docking simulation was set up by setting receptor to "unselected atoms" and ligand to "selected atoms". The MDB file containing the processed ligands to be docked were loaded. Ligand placement and refinement were set using the Alpha PMI and induced fit methods, 30 and 3 poses respectively. After the docking simulation was completed, the docking poses were scored using the S-score value (Free Energy of Binding (kcal/mol), GBVI/WSA dG), RMSD (Å) value (deviation between docking poses) and by comparison with the IMP-1088 binding mode in the ZtNMT homology model. The resulting MDB file contains the atomic coordinates for all docked inhibitors in their respective binding pockets and was used to generate an SAR report to help rationalize *in vitro* and *in vivo* activity.

**Label-free proteomics for the identification of myristoylated protein.** Myristoylated proteome enrichment for label free proteomics was performed as follows. 50 µl of NeutrAvidin® Agarose Resin slurry from Thermo were washed with 500 µl of 0.2% SDS in PBS solution three times by vortexing for 1 min and centrifugation in a microcentrifuge for 1min. 1 ml of 1mg/ml protein solution from AzRB click chemistry reaction was added to the beads and they were stirred at room temperature for 2 h. The supernatant was kept, to check enrichment efficiency. The beads were washed in three times with 500 µl 1% SDS in PBS, followed by two times with 4 M urea in 50 mM ammonium bicarbonate, followed by three times with 50 mM ammonium bicarbonate (AMBIC). The beads were resuspended in 50 µl 50 mM AMBIC and 5 µl of 100 mM TCEP and were stirred for 1 h at 37 $^{0}$C in a thermomixer (Eppendorf). The beads were pelleted in a microcentrifuge and the supernatant was swapped for another 50 µl of 50 mM AMBIC and 5 µl of 100 mM iodoacetamide left in a dark without stirring for 30 min. The beads were washed twice with 500 µl 50 mM AMBIC. 50 µl of 50 mM AMBIC was again added to the beads followed by 1 µl of trypsin solution (a vial with 20 µg Sequencing Grade Modified Trypsin from Promega dissolved in 100 µl 50 mM AMBIC) and left shaking in a thermomixer at 37 $^{0}$C overnight. The supernatant was collected and the beads resuspended in 70 µl 50 mM AMBIC and stirred for 15 min. The supernatant was collected and combined with a previous fraction. The beads were resuspended in 70 µl 1.5% TFA in MilliQ water and stirred for 15 minutes. The supernatant was again collected and combined with previous fractions. The resulting peptides solution was then desalted on a STAGE (stop and go extraction) tip.

A stack of three Empore™ SDB XC discs from Sigma was cut and fit into a p200 pipette tip. The tips were activated with 150 µl of methanol by centrifugation for 2 min at 2000g and washed with 150 µl of MilliQ water for 2 min at 2000g. The peptide samples were loaded onto the tips and centrifuged for 2 min at 2000g. The tips were washed with 150 µl of MilliQ water for 2 min at 2000g. The peptides were eluted from the tips using 60 µl of 79% acetonitrile in water by centrifugation for 2 min at 2000g. The solutions were then dried in a Thermo Savant SPD1010 speedvac and stored at -80 $^{0}$C.

**TMT-based proteomics analysis of myristate enriched proteins after NMT inhibition.** Myristoylated proteome enrichment for TMT labelled proteomics was performed as follows. 50 µl of NeutrAvidin® Agarose Resin slurry from Thermo were washed with 500 µl of 0.2% SDS in PBS solution three times by vortexing for 1 min and centrifugation in a microcentrifuge for 1min. 1 ml of 1mg/ml protein solution from AzRB click chemistry reaction was added to the beads and they were stirred at room temperature for 2 h. The supernatant was kept, to check enrichment efficiency. The beads were washed in three times with 500 µl 1% SDS in PBS, followed by two times with 4 M urea in 50 mM triethylammonium bicarbonate, followed by three times with 100 mM triethylammonium bicarbonate (TEAB). The beads were resuspended in 50 µl 100 mM AMBIC and 5 µl of 100 mM TCEP and were stirred for 1 h at 37 $^{0}$C in a thermomixer (Eppendorf). The beads were pelleted in a microcentrifuge and the supernatant was swapped for another 50 µl of 100 mM TEAB and 5 µl of 100 mM iodoacetamide left in a dark without stirring for 30 min. The beads were washed twice with 500 µl 100 mM TEAB. 30 µl of 100 mM TEAB was again added to the beads followed by 1uL of trypsin solution (a vial with 20 µg Sequencing Grade Modified Trypsin from Promega dissolved in 100 µl 100 mM TEAB) and left shaking in a thermomixer at 370C overnight. The supernatant (30 µl) was collected and the beads resuspended in 70 µl 100 mM TEAB and stirred for 15 min. The supernatant was collected and combined with the previous fraction. A quarter of the resulting solution (30 µl) was used for TMT labelling.

0.8 mg of each of the TMTsixplex™ Isobaric labels was dissolved in 41 µl anhydrous acetonitrile and then 8 µl of the solution of a reagent was added to 30 µl of corresponding peptide solution to start the labelling reaction. The solution was stirred at room temperature for 1.5 h. Then 2 µl of 5% hydroxylamine solution was added to quench the reaction and the solution was stirred for additional 15 minutes. The labelled solutions were pooled together into six-plexes and desalted on a STAGE (stop and go extraction) tip.

During the STAGE tip fractionation, some mistakes which do not undercut the results were made, it is, therefore, recommended to use a similar protocol for STAGE tip fractionation, described under "Whole lysate TMT labelled proteomics".

A stack of three Empore™ SDB RPS discs from Sigma was cut and fit into a p200 pipette tip. The tips were washed with 150 µl of methanol and then with 150 µl of MilliQ water by centrifugation for 2 min at 2000g. The peptide samples were loaded onto the tips and centrifuged for 2 min at 2000g. The tips were washed with 60 µl of 0.2% v/v trifluoroacetic acid in MilliQ water for 2 min at 2000g. Combined loading and wash flow through was saved as the first fraction. The second fraction was obtained by eluting the tip with 60 µl of 100 mM ammonium formate, 40 % v/v acetonitrile, 0.5% v/v formic acid for 2 min at 2000g. The third fraction was obtained by eluting the tip with 60 µl of 150 mM ammonium formate, 60 % v/v acetonitrile, 0.5% v/v formic acid for 2 min at 2000g. The fourth fraction was obtained by eluting the tip with 60 µl of 5% v/v ammonium hydroxide, 80 % v/v acetonitrile for 2 min at 2000g. The first fraction was desalted again. The tips made with a stack of three Empore™ SDB XC discs were activated with 150 µl of methanol by centrifugation for 2 min at 2000g and washed with 150 µl of MilliQ water for 2 min at 2000g. The flow through with the first fraction was loaded onto the tips and centrifuged for 2 min at 2000g. The tips were washed with 150 µl of MilliQ water for 2 min at 2000g. The peptides were eluted from the tips using 60 µl of 79% acetonitrile in water by centrifugation for 2 min at 2000g. All the fractions were then dried in a Thermo Savant SPD1010 speedvac and stored at -80 $^0$C.

**TMT-based proteomics analysis of the whole lysate after NMT inhibition.** Whole lysate TMT labelled proteomics was performed as follows. 1 µl 200 mM TCEP was added to 50 µl of 1mg/ml protein samples and stirred in a thermoshaker at 55 °C for 1 hr. Then 200 µl of 8 M urea in 0.1 M Tris/HCl pH 8.5 (urea solution) were added. 50 µl of protein solutions (a fifth) were transferred to a 10kDa MWCO spin column and spun 14000g for 15 min at room temperature. 200 µl 8 M urea solution was added to the filters and the samples were washed by centrifugation at 14000g for 15 min. 100 µl 0.05 M iodoacetamide in 8M urea solution was added to the filters and the samples were left in the dark for 30 min at room temperature. The samples were centrifuged for 15 min at 14000g and washed two times with 100 µl 8 M urea solution. The samples were then washed three times with 100 µl 100 mM TEAB solution. 50 µl 100 mM TEAB and 1 µl of trypsin (a vial with 20 µg Sequencing Grade Modified Trypsin from Promega dissolved in 100 µl 100 mM TEAB) were added to the filters and they were left stirring in a thermoshaker at 37 °C overnight.

TMTsixplex™ Isobaric labels were dissolved in 41 µl anhydrous acetonitrile. 10 µl of each label was transferred to their respective filters and they were stirred at room temperature for 1 h. 2 µl of 5% hydroxylamine was added to the samples and stirred at room temperature for 30 min. Peptides were eluted from the spin columns by two washes with 40 µl 100 mM TEAB solution and one wash with 50 µl 0.5 M NaCl solution. Samples were then combined into six-plexes and partially dried in a Speedvac to a volume of around 200 µl to remove acetonitrile. 2 µl of trifluoroacetic acid was added before STAGE tip fractionation.

A stack of three Empore™ SDB RPS discs from Sigma was cut and fit into a p200 pipette tip. The peptide samples were loaded onto the tips and centrifuged for 2 min at 2000g. The tips were washed with 60 µl of 0.2% v/v trifluoroacetic acid in MilliQ water for 2 min at 2000g. The first fraction was obtained by eluting the tip with 60 µl of 100 mM ammonium formate, 40 % v/v acetonitrile, 0.5% v/v formic acid for 2 min at 2000g. The second fraction was obtained by eluting the tip with 60 µl of 150 mM ammonium formate, 60 % v/v acetonitrile, 0.5% v/v formic acid for 2 min at 2000g. The third fraction was obtained by eluting the tip with 60 µl of 5% v/v ammonium hydroxide, 80 % v/v acetonitrile for 2 min at 2000g. All the fractions were then dried in a Thermo Savant SPD1010 speedvac and stored at -80 $^0$C.

**LC-MS/MS label-free (LFQ) and TMT methods.** Dried peptide fractions were dissolved in 15 µl of LC-MS grade $H_2O$ containing 2% (v/v) acetonitrile and 0.5% (v/v) TFA. 3 µl of the solution were injected into the LC-MS/MS system. Peptides were separated on an Acclaim PepMap RSLC column 50 cm × 75 µm inner diameter (Thermo Fisher Scientific) using a 2 h (LFQ) or 3 h (TMT) acetonitrile gradient in 0.1% aqueous formic acid at a flow rate of 250 nl/min. Easy nLC-1000 was coupled to a QExactive mass spectrometer via an easy-spray source (all Thermo Fisher Scientific). The QExactive was operated in data-dependent mode with survey scans acquired at a resolution of 70,000 at m/z 200. Scans were acquired from 350 to 1800 m/z. Up to 10 of the most abundant isotope patterns with charge +2 or higher from the survey scan were selected with an isolation window of 3.0 m/z (LFQ) or 1.6 m/z (TMT) and fragmented by HCD with normalized collision energies of 25 W (LFQ) or 31 W (TMT). For LFQ, maximum ion injection times for the survey scan and the MS/MS scans (acquired with a resolution of 17,500 at m/z 200) were 250 and 80 ms, respectively; the ion target value for MS was set to 106 and for MS/MS to 105. For TMT, the maximum ion injection times for the survey scan and the MS/MS scans (acquired with a resolution of 35,000 at m/z 200) were 20 and 120 ms, respectively; the ion target value for MS was set to 106 and for MS/MS to $2\times10^{5}$, and the intensity threshold was set to $1.7\times10^{3}$.

Proteomics results (*.RAW files) were processed using MaxQuant 1.5.5.1 or PEAKS 8.0 using latest releases of *Zymoseptoria tritici* proteome at uniprot.org.

In MaxQuant, digestion was set as Trypsin/P, oxidation (M) and acetylation (N-term) were set as variable modifications, carbamidomethylation (C) was set as fixed modification and other settings were as by default.

In PEAKS, digestion was set as Trypsin, precursor mass tolerance was set at 5.0 ppm, fragment ion mass tolerance was set 0.1 Da. Oxidation (M) and acetylation (N-term) were set as variable modifications, carbamidomethylation (C) was set as fixed modification and PEAKS PTM was used to find additional modifications. Additional variable modification on glycines at *N*-termini with mass 463.2907 was used to find MS/MS spectra of peptides carrying the alkyne modified myristoylated analogue (YnMyr) clicked with AzRB capture reagent.

Perseus 1.5.5.3 was used to analyze file proteinGroups.txt obtained as result of MaxQuant search. Potential contaminants, reverse and only identified by site proteins were discarded before the analysis. All data was transformed using log2(x) function before any further manipulation. LFQ data was normalized by subtracting a median of each data column. TMT data was normalized by first subtracting a median from each data column, followed by subtracting a mean of six values of a TMT six-plex for each such six-plex. The data was filtered to retain only those proteins that have at least two measurements across replicates for every experimental condition. *t*-test was performed using default Perseus settings, volcano plot representations have a visual cut-off line with parameters: s0=0.1, FDR=0.05.

Hierarchical clustering was performed by Pearson correlation. In Fisher's exact test, the default Benjamini-Hochberg FDR value of 0.02 was used for significance cut-off.

**Table S1. Potency of inhibitors tested in enzymatic CPM ($IC_{50}$) and viability MTS ($EC_{50}$) assays, relation between them (ratio $EC_{50}/IC_{50}$) and the results of MOE docking (S-Score, kcal/mol and RMSD, Å).**

| Inhibitor name (reference) | Inhibitor structure | $IC_{50}$ (μM) | $IC_{50}$ CI (μM) | $EC_{50}$ (μM) | $EC_{50}$ CI (μM) | $EC_{50}/IC_{50}$ | S-score, kcal/mol | RMSD, Å |
|---|---|---|---|---|---|---|---|---|
| **IMP-1031**[2] | 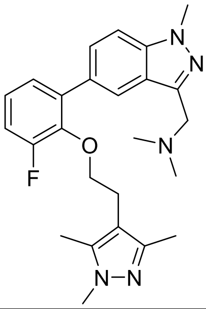 | 0.016 | 0.0092-0.017 | 51 | 46-58 | 4400 | -8.71 | 1.61 |
| **IMP-1034**[1] | 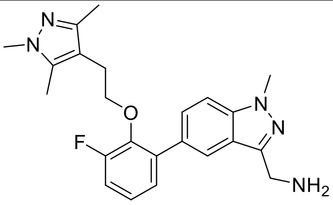 | 0.032 | 0.017-0.056 | | | | -8.35 | 1.02 |
| **IMP-1036**[1] | 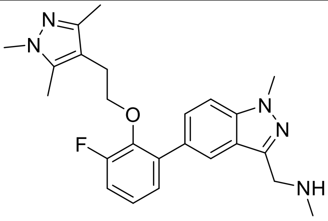 | 0.030 | 0.016-0.053 | | | | -8.43 | 1.00 |
| **IMP-1088**[1,2] | 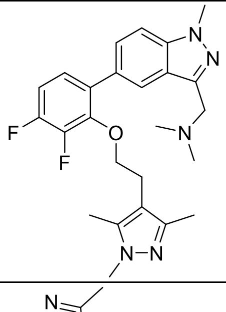 | 0.014 | 0.011-0.023 | 23 | 22-25 | 2100 | -8.66 | 1.45 |
| **IMP-1089**[1] | 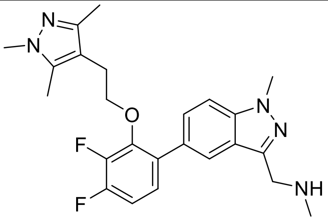 | 0.0067 | 0-0.013 | 46 | 39-52 | 7800 | -8.47 | 1.17 |
| **IMP-1090**[1] | 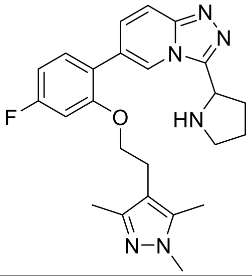 | 22 | 2.5-1000 | | | | -8.53 | 1.03 |

| | | | | | | | | |
|---|---|---|---|---|---|---|---|---|
| **IMP-1091**[1] | 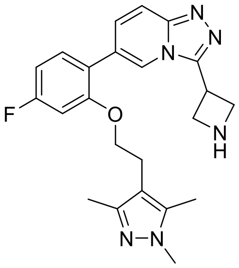 | 2.1 | 1.2-1000 | | | | -8.15 | 1.59 |
| **IMP-1093**[1] | 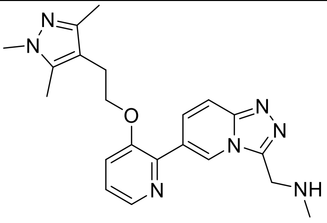 | >300 | | | | | -8.13 | 1.32 |
| **IMP-1094**[1] | 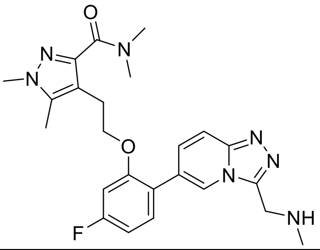 | 0.086 | 0.06-0.13 | | | | -8.79 | 0.897 |
| **IMP-1097**[1] | 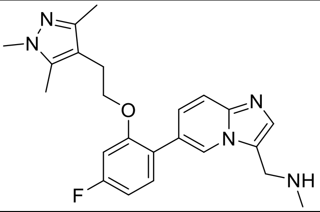 | 0.20 | 0.02-0.60 | | | | -8.2 | 1.59 |
| **IMP-1100**[1] | 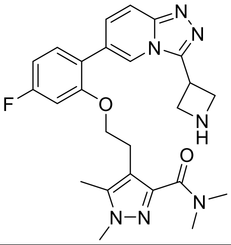 | 0.016 | 0.009-0.025 | >500 | | | -9.19 | 0.959 |
| **IMP-1111**[1] | 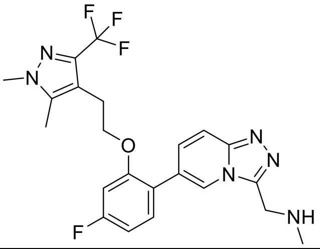 | 0.70 | 0.50-0.96 | | | | -8.38 | 1.59 |
| **IMP-1112**[1] | 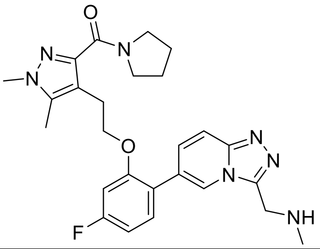 | 0.42 | 0.25-0.77 | | | | -9.28 | 1.57 |

| | | | | | | | | |
|---|---|---|---|---|---|---|---|---|
| **IMP-1124**[1] | 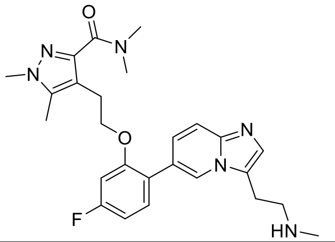 | 0.013 | 0-0.030 | | | | -9.46 | 1.47 |
| **IMP-1131**[1] | 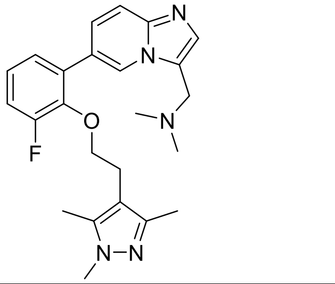 | 2.4 | 1.6-4.2 | | | | -8.25 | 0.986 |
| **IMP-1134**[1] | 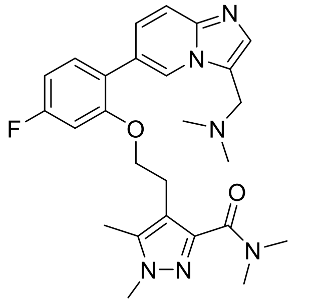 | <0.001 | | 27 | 23-32 | >32000 | -9.42 | 1.00 |
| **IMP-1179**[1] | 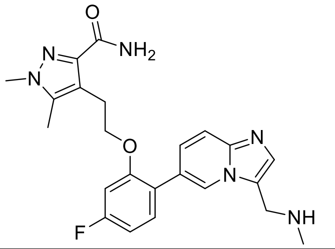 | 0.28 | 0.20-0.41 | | | | -8.52 | 1.44 |
| **IMP-1185**[1] | 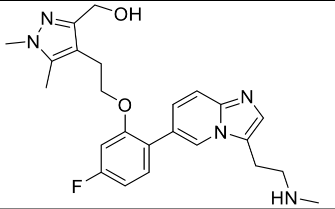 | 4.0 | 2.5-9.4 | | | | -8.38 | 1.17 |
| **IMP-1195**[1] | 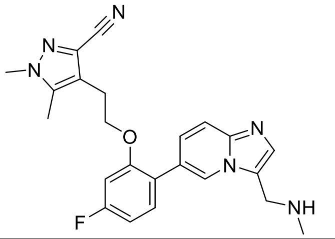 | <0.001 | 0-0.020 | | | | -8.24 | 1.25 |
| **IMP-162**[3] | 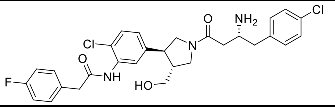 | 1.9 | 1.6-2.2 | 18 | 14-24 | 13 | | |
| **IMP-228**[4] | 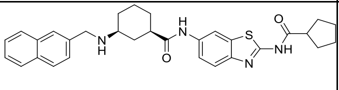 | >300 | | | | | | |

| | | | | | | | | |
|---|---|---|---|---|---|---|---|---|
| **IMP-277**[5] | 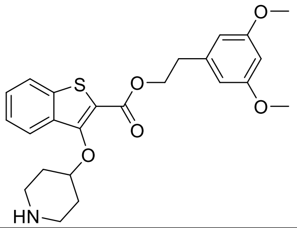 | >300 | | | | | | |
| **IMP-329**[6] | 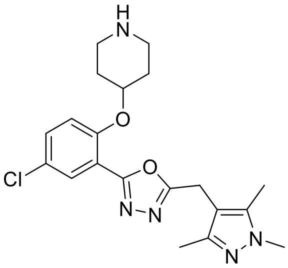 | 0.87 | 0.72-1.1 | 280 | 210-370 | 430 | | |
| **IMP-366**[7] | 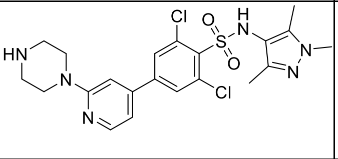 | 0.050 | 0.043-0.057 | 110 | 74-210 | 4100 | | |
| **IMP-917**[2] | 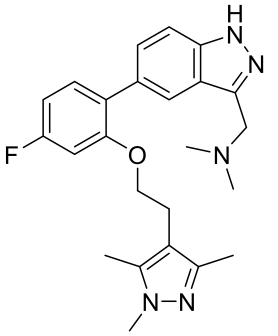 | 0.16 | 0.084-0.31 | | | | -8.22 | 1.22 |
| **IMP-964**[6] | 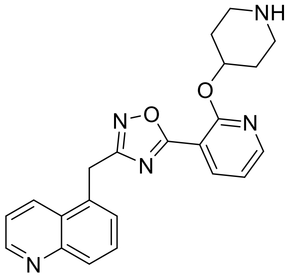 | 0.52 | 0.10-2.5 | 160 | 110-220 | 420 | | |
| **IMP-970**[5] | 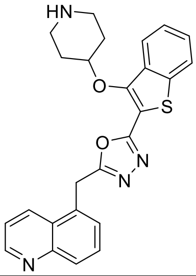 | 2.8 | 1.4-5.9 | | | | | |

### Supporting Information References
